# High-speed three-dimensional random access scanning with SPARCLS

**DOI:** 10.1101/2024.07.08.602445

**Authors:** Caroline Berlage, Urs L. Böhm, Ana Sanchez Moreno, Julia Ledderose, Albert Gidon, Matthew E. Larkum, Andrew Plested, Benjamin Judkewitz

## Abstract

High-speed volumetric imaging is crucial for observing fast and distributed processes such as neuronal activity. Multiphoton microscopy helps to mitigate scattering effects inside tissue, but the standard raster scanning approach limits achievable volume rates. Random-access scanning can lead to a considerable speed-up by sampling only pre-selected locations, but existing techniques based on acousto-optic deflectors are still limited to a point rate of up to ∼50 kHz. This limits the number of parallel targets at the high acquisition rates necessary, for example, in voltage imaging or imaging of fast synaptic events.

Here we introduce SPARCLS, a method for 3D random-access scanning at up to 340 kHz point rate using a single 1D phase modulator. We show the potential of this method by imaging synaptic events with fluorescent glutamate sensors in mammalian organotypic slices as well as in zebrafish larvae.

## 1. Introduction

Optical microscopy plays a crucial role in biomedical research. For imaging living tissues, one of the most widespread techniques is two-photon (2P) laser scanning microscopy, because it enables high-resolution imaging at depth: Fluorescence is excited nonlinearly in the vicinity of a moving focus, thus reducing the susceptibility to scattering [1, 2].

The imaging speed of such laser scanning techniques, however, is limited by the inertia of scanning mirrors for lateral scanning, and even more by mechanisms for axial scanning [3]. Fundamentally, imaging rate is also limited by the fluorescence lifetime, which restricts the pixel dwell time to a minimum on the order of several nanoseconds. Especially for volumetric imaging, this severely limits the rate at which dynamic processes can be observed.

When observing biological activity, particularly in neuroscience [4–8], structures are typically sparse and not all points within the field of view (FOV) are equally relevant. Random access microscopy [9] takes advantage of this fact by targeted sampling of pre-selected locations (Fig. 1a). As long as a scanning mechanism with negligible inertia is used, this enables a considerable speed-up, but conventional galvanometric scanning mirrors do not fit this criterion.

**Fig. 1.**
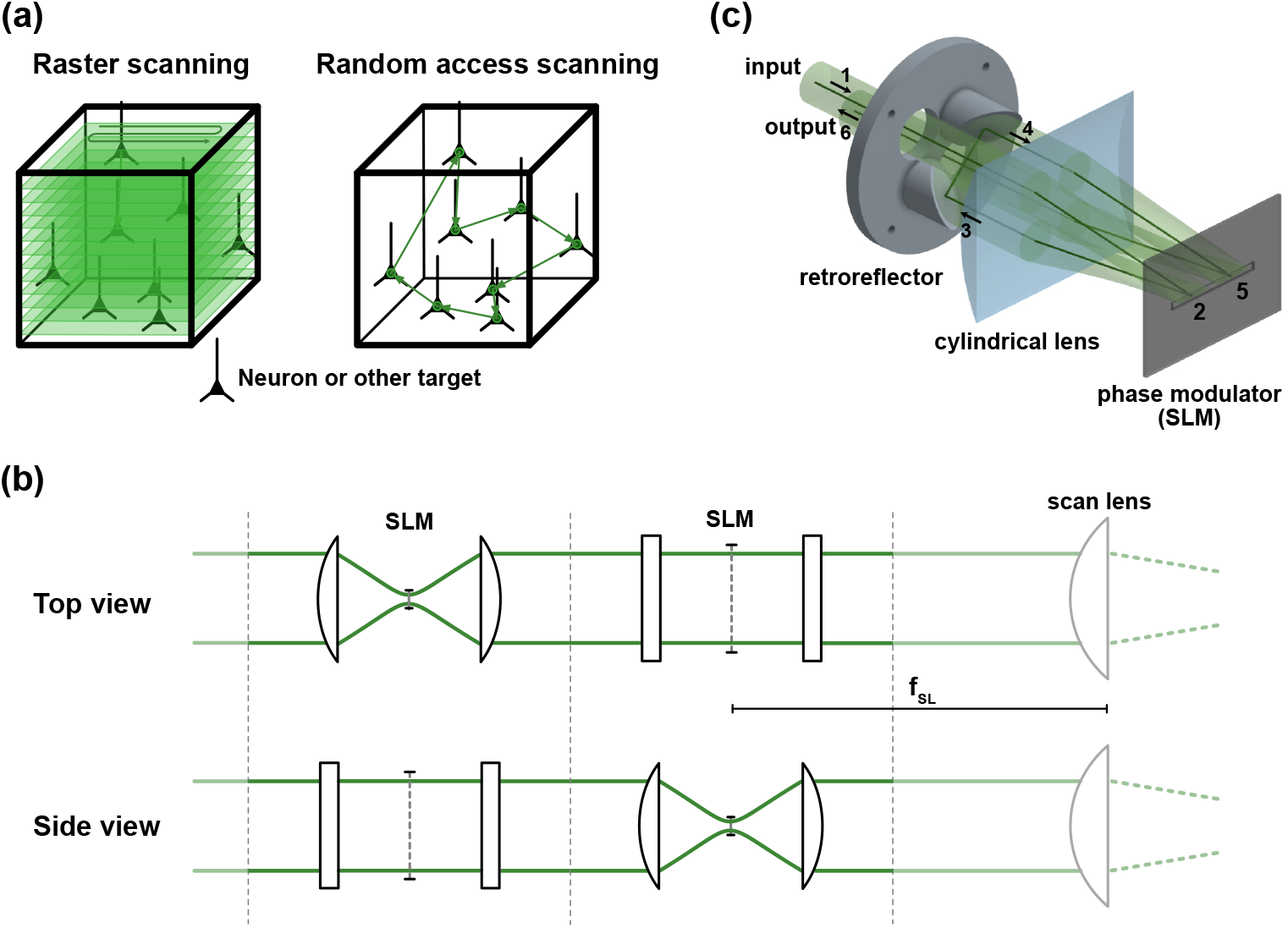
Three-dimensional random-access imaging using a single 1D spatial light modulator. (a) Raster scanning a focused beam across the sample generates a detailed 3D image, but volume acquisition rates are limited by inertia and by the fluorescence lifetime. Random-access scanning allows faster volume rates by sampling only pre-selected locations. (b) Unfolded setup to illustrate the beam path. A collimated beam enters the module. The cylindrical lens creates a line focus on the SLM, where the first axis is modulated. The beam is recollimated and then focused orthogonal to the first axis, modulated on the SLM, and recollimated. A scan lens and tube lens (not shown) image the SLM plane onto the objective’s back aperture. (c) 3D visualization of the compact scanning module. The beam enters through a 12.5 mm diameter input port on the top left quadrant (1) and is first modulated on the front half of the SLM (2). The custom retroreflector rotates the beam at 90 degrees (3), so the same cylindrical lens and SLM can be reused in the back half (4-5). The modulated beam exits the module at the bottom right (6). This module can be added to a conventional multiphoton microscope.

Non-mechanical custom beam steering has been realized at tens of kHz update rate with spatial light modulators (SLMs) [10–12] or digital micromirror devices (DMDs) [13]. The most established non-mechanical approach uses acousto-optic deflectors (AODs) and enables scanning at 35-50 kHz in a single plane [9, 14, 15] and in three dimensions [16–19]. Synchronization with laser pulse repetition rates or counterpropagating nonlinearly chirped waves can also enable beam shaping and wavefront corrections [7, 20, 21] as well as optogenetic stimulation [22]. Drawbacks of AOD-based scanning are the highly dispersive crystal material, high setup complexity and high costs, as most setups require at least 3-4, some even 6 AODs [21], controlled with analogue electronics. Additionally, it would be preferable to further increase the random access scanning rate, as kilohertz acquisition rates necessary for voltage or neurotransmitter imaging still limit the available number of targets. However, the time required for an acoustic wave to cross an AOD aperture limits practically attainable random access rates to ≤50 kHz.

Here, we present SPARCLS (scanning points and rapid custom access using a linear SLM): a method for 3D random-access scanning at up to 340 kHz using a MEMS-based 1D spatial light modulator in a compact and modular optical setup that can be used in conjunction with existing multiphoton microscopes. In addition to raising the speed of random access point scanning by nearly an order of magnitude, we demonstrate the method’s potential by volumetrically recording fast synaptic events in organotypic slices and zebrafish larvae at nearly 100 sites simultaneously.

## 2. Results

### 2.1. Principle and Characterization

Recent work on 1D beam steering and wavefront shaping with reflective MEMS-based linear SLMs has demonstrated astonishing update rates of up to 350 kHz [23–25]. We asked whether such devices could be used as a basis for ultrafast 3D scanning.

By using two linear modulators in a crossed configuration, it should in principle be possible to create any linearly separable 2D pattern. This includes both lateral deflections and defocus (to access the third dimension). Additionally, many Zernike polynomials can be closely approximated by linearly separable patterns to correct for aberrations [20]. To illustrate the principle of 3D modulation using linear modulators, an unfolded setup using two (hypothetically transmissive) linear SLMs and four cylindrical lenses is shown in Fig. 1b. In short, a cylindrical lens focuses the initially collimated beam to a line on the first SLM, where it is phase modulated along the first dimension. A second cylindrical lens recollimates the beam before the same process occurs rotated by 90 degrees. Using a scan lens and tube lens, the SLM plane is then imaged onto the back focal plane of a microscope objective.

Considering this basic design and hardware, we identified several key hurdles: First, as a beam deflection by the first SLM will shift the second line focus up or down, the short axis of the second SLMs active area will limit the scanning range of the first SLM. This necessitates a characterisation of effective pixel size along this axis and the use of short focal length lenses. As currently available phase modulators are reflective, short focal lengths make it difficult to separate input and output. The second hurdle is to adequately synchronize the SLMs, ideally at a multiple of their refresh rate (i.e. ≥1 MHz). Finally, the only available device for high-speed scanning at hundreds of kHz (Silicon Light Machines F1088-P) is designed for a full stroke at blue and UV wavelengths, falling far short of full phase modulation at the near-infrared (NIR) wavelengths used for biological multiphoton imaging.

We addressed the first two hurdles with the design shown in Figure 1c. We solved the synchronization challenge by using the same SLM device twice, while rotating the beam between both passes. This also allowed us to minimize focal lengths and make a compact optical module: a collimated, approximately 10 mm-diameter beam enters through an input port in the top left quadrant of a custom mount. The cylindrical lens creates a line focus on the left half of the phase modulator, which is then reflected and recollimated in the lower left quadrant before reaching the retroreflector. Two elliptical mirrors mounted on the retroreflector reflect the beam back towards the top right quadrant of the cylindrical lens and, at the same time, rotate the beam by 90 degrees, so that the second modulation on the phase modulator is orthogonal to the first. The beam then exits the module through an output port in the bottom right quadrant. The SLM plane is imaged onto the back focal plane of the objective with unit magnification using two achromatic lenses as scan and tube lens. Unmodulated light and unwanted higher diffraction orders are blocked by a rectangular absorptive aperture at the scan lens focal plane, while the first order is used for scanning. More information on the microscope setup can be found in the Supplementary Information.

When calibrating the relationship between command signal and optical phase retardance (see Supplementary Information), we determined the effective maximum optical path length stroke to be ∼400 nm, in agreement with its design for blue wavelengths. This corresponds to a phase stroke of ∼0.8π at a wavelength of 940 nm (at the incidence angle used in our setup), and may at first sight rule out any use with NIR light. However, even limited stroke leads to noticeable diffraction efficiency, which we calculated to be nearly 50% of its maximum value (see Supplementary Figure S2). Due to the combined effects of diffraction efficiency, two reflections and phase modulations, as well as transmission losses, the overall power efficiency of the scan unit is approximately 5%. However, given that commercially available fs-pulsed light sources output powers of several Watts, this efficiency was sufficient even for *in vivo* experiments (see Applications, below).

We determined the FOV for a 25x objective (Leica HC Fluotar L 25x/0.95 W VISIR; effective NA 0.6) by imaging fluorescent beads dispersed in agarose (Figure 2a; 1 μm yellow-green FluoSpheres, ThermoFisher). As previously discussed, the scan range along the short lateral axis is limited by the height of the pixels. While the reported usable height (with guaranteed phase stroke) is only 75 μm, we found that the ribbon-based MEMS pixels have a much larger usable height of up to 160 μm. The FOV using a 40x/0.8 NA objective is shown in Supplementary Figure S4. We used a modal adaptive optics algorithm [26] based on fluorescence intensity to correct for linearly separable system aberrations (see Supplementary Information and Supplementary Figure S1). For both objectives, the point spread function after system correction is shown in Supplementary Figure S3.

**Fig. 2.**
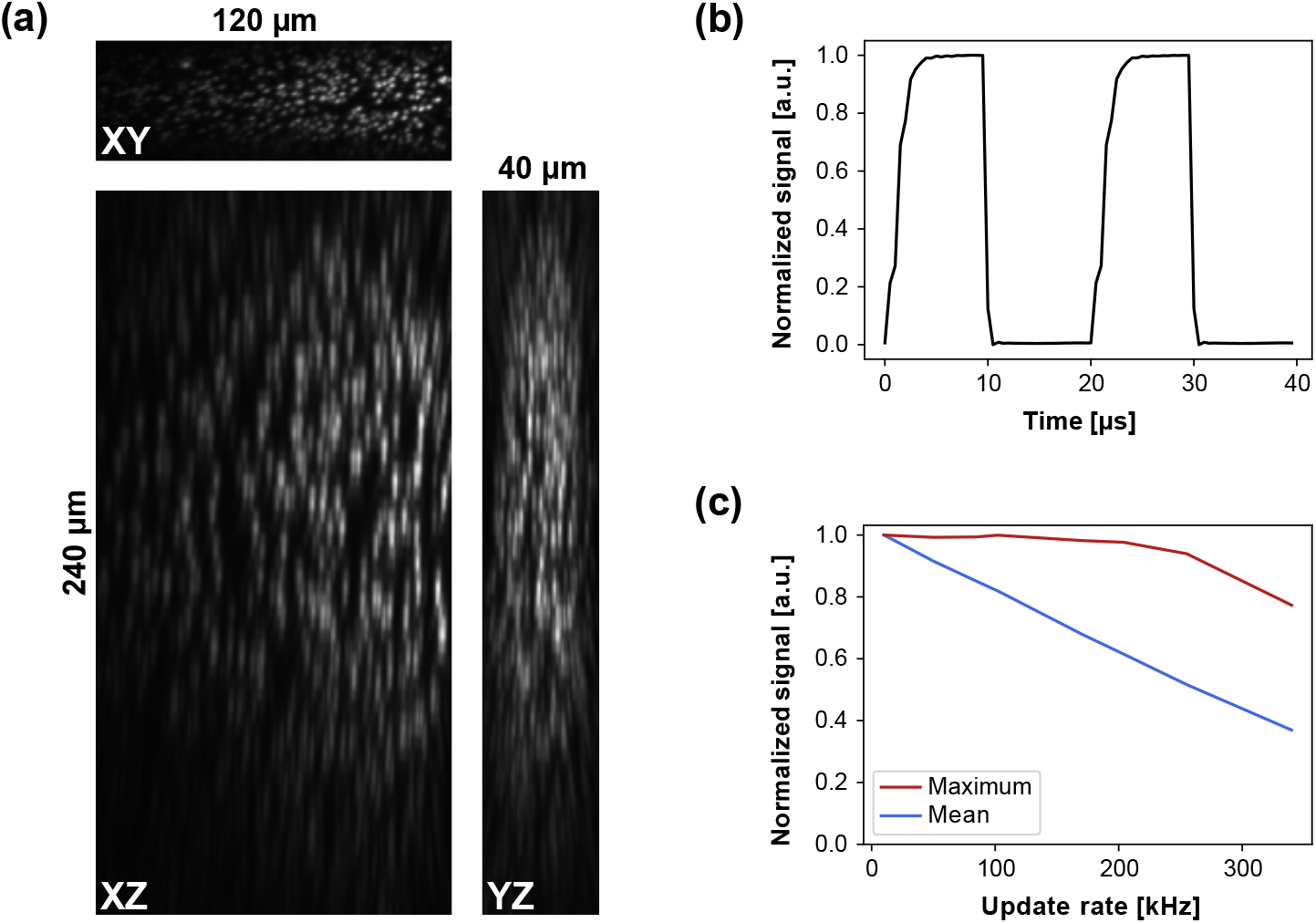
Spatial and temporal characterization. (a) Three-dimensional FOV visualized with 1 μm diameter beads using a 25x objective with effective NA 0.63. (b) Temporal switching characteristics: The phase pattern switches between a bright and dark target within the FOV. The phase modulator is updated at 100 kHz and the PMT signal is acquired at 2 MHz. The 10-90% switching time is 2.75 μs ± 0.25 μs. (c) Bright and dark targets alternate as in (b). Both the maximum (red) and mean (blue) values of the bright target, normalized to the value at 10 kHz, are plotted at each sampling rate.

To investigate the temporal characteristics of the scan unit, we selected a bright and a dark spot within the FOV as targets. The phase modulator was set to update at 100 kHz, while the fluorescence intensity was acquired at 2 MHz. The time course of fluorescence intensity over two such cycles is shown in Fig. 2b, showing a 10-90% switching time of 2.75 μs ± 0.25 μs. To further quantify switching efficiency, we investigated the maximum and mean signal at varying SLM refresh rates. The SLM update rate was set to integer divisors of the data acquisition rate (1020 kHz), up to 340 kHz. We then determined the maximum and mean signal at the bright target normalized to the maximum intensity value at the lowest SLM refresh rate, in this case 10 kHz. For 340 kHz, the fluorescence reaches 77 % of the maximum at lower rates. The average signal decreases linearly and reaches 37% of the maximum at 340 kHz, while at 50 kHz, faster than most AOD setups, it is still at almost 92% (Fig. 2C). This means that at this speed, only 8% of the signal is lost due to switching time. In contrast, for AODs, access times on the order of 20 μs [27] do not leave any dwell time at this rate.

### 2.2. Applications

Fast 3D random access scanning is particularly suited for imaging neuronal activity. We demonstrated this by measuring glutamate release with the glutamate sensor iGluSnFR3 [28] in mouse organotypic slices and in zebrafish larvae. To resolve finer structures and increase power density at the focus, the objective was replaced by 40x/0.8 NA objective (NIR Apo 40x/0.8W, Nikon), which reduces the FOV to 25 x 75 x 120 μm^3^.

We first imaged glutamate in mouse organotypic hippocampal slice cultures (for more details, see Supplementary Information). Fig. 3a shows a raster-scanned overview stack with 5 μm Z-step size, which formed the basis for manual location selection (blue dots). Fluorescence was then recorded at these 59 locations at a pixel rate of 250 kHz. Selected traces showing the change in fluorescence dF/F0 in response to glutamate release at these locations are shown in Fig. 3b. Each trace corresponds to a single pixel without any spatial averaging. dF/F traces over four minutes show little to no bleaching (18 mW excitation power at sample). Fig. 3c shows a 30-second excerpt of the whole recording corresponding to the black boxed region in Figure 3b.

**Fig. 3.**
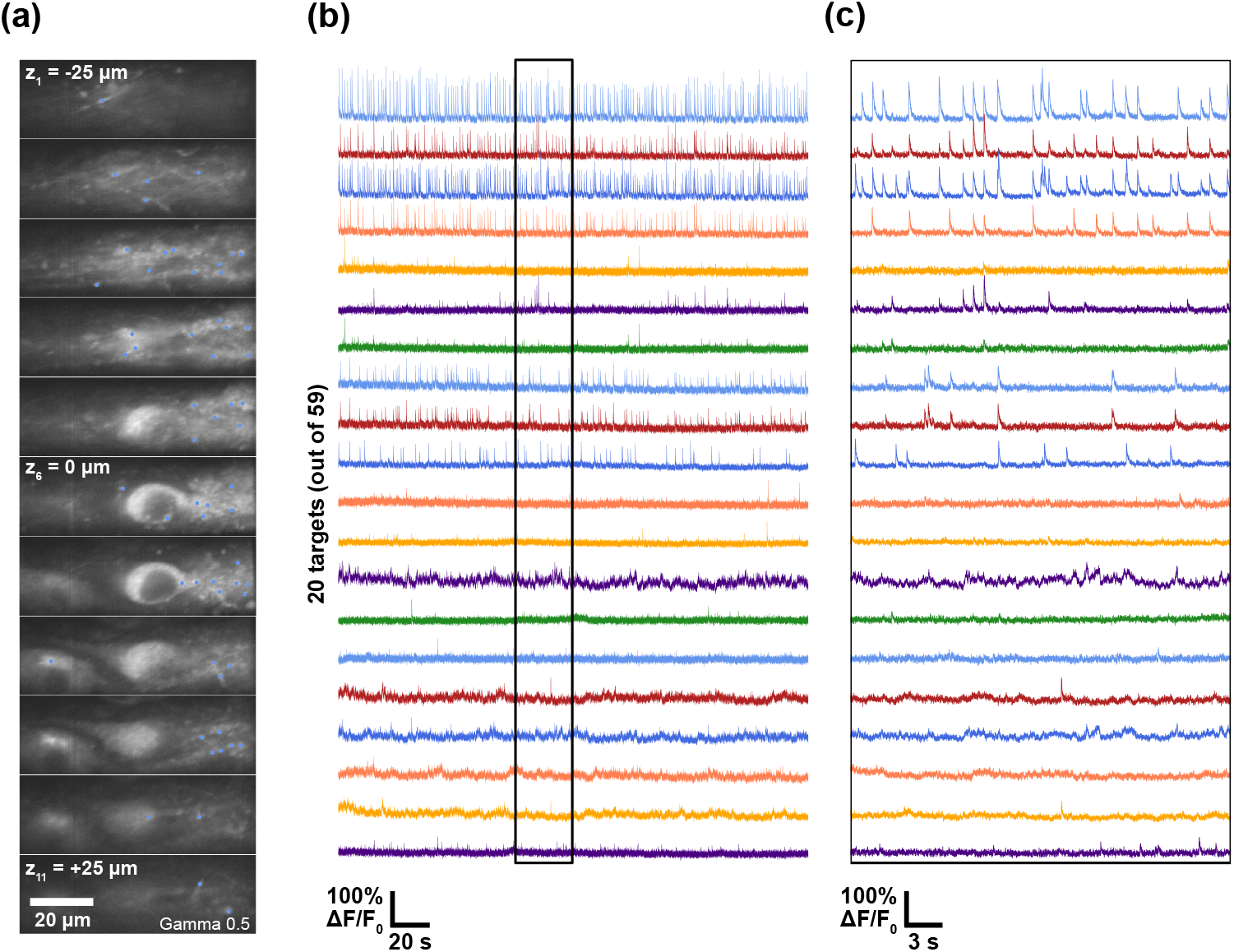
3D random access microscopy using iGluSnFR3 in mouse organotypic slices at 250 kHz point rate. (a) A raster-scanned overview stack of an organotypic slice labeled with iGluSnFR3. A Gamma correction was applied to emphasize dimmer structures (for visibility in this panel, not during analysis). Manually selected targets for random-access scanning are marked as blue dots. To increase overall activity, slices were imaged in a solution containing 20 mM KCl and 0.2 mM 4AP. (b) Change in fluorescence dF/F0 acquired at 250 kHz. For display, traces were filtered with a rolling average window of size 50. (c) Zoom of the region in (b) marked with a black rectangle.

For *in vivo* imaging of zebrafish larvae, we transiently expressed iGluSnFR3 in neurons. Fig. 4a shows a representative raster-scanned overview stack of labeled neurons in the olfactory bulb. 90 targets were manually selected and observed for 240 s (Fig. 3b) at 340 kHz point rate and 26 mW laser power. Fig. 3c shows a zoom of the highlighted region in (b).

**Fig. 4.**
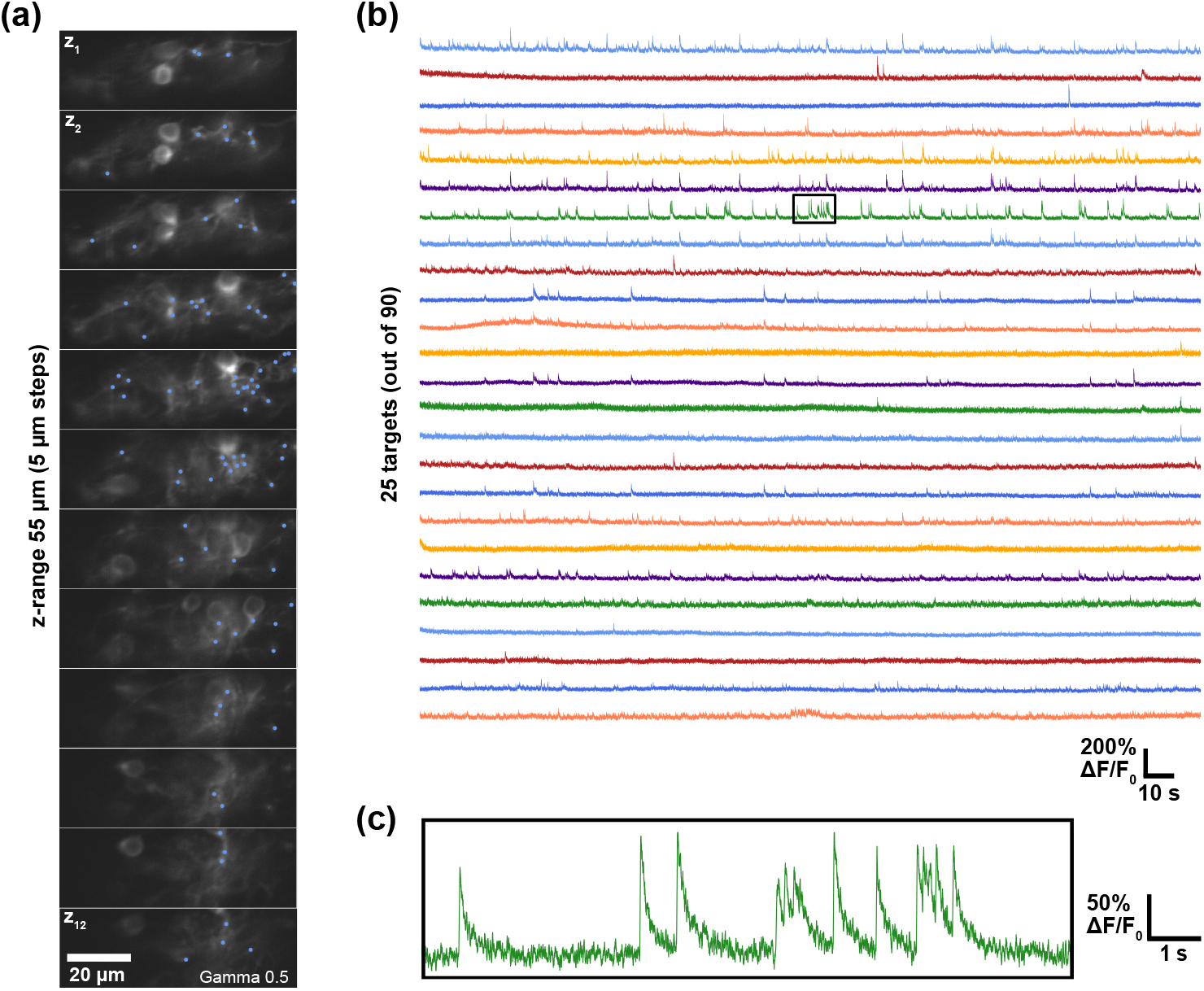
3D random access microscopy using iGluSnFR3 in larval zebrafish olfactory bulb at 340 kHz point rate. (a) Raster scanned overview stack of part of the olfactory bulb of a 3 dpf zebrafish larva. A Gamma correction was applied to emphasize dimmer structures. Manually selected targets for random-access scanning are marked as blue dots. Fish were paralyzed and overall activity was increased using pentylenetetrazol (PTZ). (b) Change in fluorescence dF/F0 acquired at 340 kHz. For display, traces were filtered with a rolling average window of size 68. (c) Zoom of the region in (b) marked with a black rectangle.

## 3. Discussion

In this work, we presented a new mechanism for 3D random-access scanning at refresh rates exceeding 300 kHz, based on a single MEMS-based 1D phase modulator in a compact and low-cost module that could be attached to existing multi-photon microscopes.

Although spatially or temporally multiplexed raster scanning methods [29–33] can now operate close to the fluorescence lifetime limit, this still imposes limits on the volume rate for large numbers of pixels. For single plane imaging of a large number of targets, line-scan tomographic methods such as scanned-line angular projection [34] combine advantages of raster scanning and random access scanning, as they allow post hoc motion correction at high frame rates. However, kHz rate scanning is limited to 2D planes.

By exclusively visiting sites of interest in rapid succession, random access scanning can reach higher scan rates in 3D, provided that the point rate is sufficiently large compared to the number of targets. Recent work [24, 35] has demonstrated that scattering media can be used to implement fast random access scanning via a 1D-to-2D transform and complex wavefront shaping. However, in complex wavefront shaping, the power in the focus is typically orders of magnitude smaller than the power deposited to the rest of the sample [36], which makes in vivo imaging challenging. Compared to AOD-based scanning [37], the standard for random-access scanning, SPARCLS improves the point rate by nearly an order of magnitude, although currently at a smaller FOV.

The FOV of our scanning approach is limited by two factors. First, the number of SLM pixels which limit the number of different scan positions, and second, clipping on the edges of the linear phase modulator on the second reflection. In addition to increasing pixel number, it might be possible to manufacture taller pixels for a larger FOV along the short lateral dimension. Beyond this, combining random access scanning with galvo scanning, i.e. shifting the center of the SLM scanning range, would enhance the accessible FOV.

Similar to all random access scanning methods, we require prior knowledge of the sample. Thus, a raster scan is performed before target selection. As this is not continuously updated, random access scanning is susceptible to motion artifacts. However, work using AODs has shown that it is possible to perform random access scanning even in awake, behaving animals using patch scanning [19] or PSF shaping [7, 22], which could also be possible using this SLM-based scanning mechanism. The faster refresh rate compared to AODs, as well as real-time control of the phase patterns [24], could also make closed-loop motion correction [38] possible.

Except for a 3D-printed retroreflector mount, the module uses only off-the-shelf components. In the future, its performance could be optimized by using custom optical elements. First, the acylindrical lens optimized for 780 nm could be replaced by an achromatic cylindrical lens designed for the wavelength range around 900-940 nm, where many indicators can be excited. This would reduce chromatic aberrations as well as allow a shorter focal length, thus increasing the FOV. Second, the phase modulator is designed for visible light. A modulator with a higher phase stroke would improve the diffraction efficiency and thus the transmission of the scan unit by a factor of up to two (see Figure S2), leading to a four-fold improvement due to the two reflections.

We used a modal AO algorithm to correct for system aberrations in the setup. In principle, the same mechanism could be used to also correct low order sample aberrations such as the zebrafish head curvature. It would be possible to apply individual correction patterns for every target point. Random access scanning using AODs is continuously finding wider application as faster and brighter fluorescent indicators and optogenetic actuators emerge. The approach presented here allowed us to increase the point scan rate of random access scanning by nearly an order of magnitude. The method enabled us to image iGluSnFR3, as one example of fast indicators of neuronal activity. In the future, it could be used for fast calcium imaging, 2P voltage imaging or for 3D patterned photostimulation in conjunction with optogenetic actuators.

## Acknowledgments

We thank Spencer Smith and Maximilian Hoffmann for discussions and Marc Renz for technical assistance with screening of zebrafish.

## References

1. W. Denk, J. H. Strickler, and W. W. Webb, “Two-Photon Laser Scanning Fluorescence Microscopy,” Science 248, 73–76 (1990).

2. F. Helmchen and W. Denk, “Deep tissue two-photon microscopy,” Nat. Methods 2, 932–940 (2005).

3. J. Wu, N. Ji, and K. K. Tsia, “Speed scaling in multiphoton fluorescence microscopy,” Nat. Photonics 15, 800–812 (2021).

4. T.-W. Chen, T. J. Wardill, Y. Sun, et al., “Ultrasensitive fluorescent proteins for imaging neuronal activity,” Nature 499, 295–300 (2013).

5. A. S. Abdelfattah, T. Kawashima, A. Singh, et al., “Bright and photostable chemigenetic indicators for extended in vivo voltage imaging,” Science 365, 699–704 (2019).

6. S. Chamberland, H. H. Yang, M. M. Pan, et al., “Fast two-photon imaging of subcellular voltage dynamics in neuronal tissue with genetically encoded indicators,” eLife 6, e25690 (2017).

7. V. Villette, M. Chavarha, I. K. Dimov, et al., “Ultrafast Two-Photon Imaging of a High-Gain Voltage Indicator in Awake Behaving Mice,” Cell 179, 1590–1608.e23 (2019).

8. Z. Liu, X. Lu, V. Villette, et al., “Sustained deep-tissue voltage recording using a fast indicator evolved for two-photon microscopy,” Cell 185, 3408–3425.e29 (2022).

9. A. Bullen, S. Patel, and P. Saggau, “High-speed, random-access fluorescence microscopy: I. High-resolution optical recording with voltage-sensitive dyes and ion indicators,” Biophys. J. 73, 477–491 (1997).

10. Y. Xue, L. Waller, H. Adesnik, and N. Pégard, “Three-dimensional multi-site random access photostimulation (3D-MAP),” eLife 11, e73266 (2022).

11. G. Faini, D. Tanese, C. Molinier, et al., “Ultrafast light targeting for high-throughput precise control of neuronal networks,” Nat. Commun. 14, 1888 (2023).

12. C. Telliez, V. D. Sars, V. Emiliani, and E. Ronzitti, “Descanned fast light targeting (deFLiT) two-photon optogenetics,” Biomed. Opt. Express 14, 6222–6232 (2023).

13. Q. Geng, C. Gu, J. Cheng, and S.-c. Chen, “Digital micromirror device-based two-photon microscopy for three-dimensional and random-access imaging,” Optica 4, 674 (2017).

14. R. Salomé, Y. Kremer, S. Dieudonné, et al., “Ultrafast random-access scanning in two-photon microscopy using acousto-optic deflectors,” J. Neurosci. Methods 154, 161–174 (2006).

15. V. Iyer, T. M. Hoogland, and P. Saggau, “Fast Functional Imaging of Single Neurons Using Random-Access Multiphoton (RAMP) Microscopy,” J. Neurophysiol. 95, 535–545 (2006).

16. G. D. Reddy and P. Saggau, “Fast three-dimensional laser scanning scheme using acousto-optic deflectors,” J. Biomed. Opt. 10, 064038 (2005).

17. G. D. Reddy, K. Kelleher, R. Fink, and P. Saggau, “Three-dimensional random access multiphoton microscopy for functional imaging of neuronal activity,” Nat. Neurosci. 11, 713–720 (2008).

18. G. Katona, G. Szalay, P. Maák, et al., “Fast two-photon in vivo imaging with three-dimensional random-access scanning in large tissue volumes,” Nat. Methods 9, 201–208 (2012).

19. K. M. N. S. Nadella, H. Roš, C. Baragli, et al., “Random-access scanning microscopy for 3D imaging in awake behaving animals,” Nat. Methods 13, 1001–1004 (2016).

20. W. Akemann, J.-F. Léger, C. Ventalon, et al., “Fast spatial beam shaping by acousto-optic diffraction for 3D non-linear microscopy,” Opt. Express 23, 28191 (2015).

21. G. Konstantinou, P. A. Kirkby, G. J. Evans, et al., “Dynamic wavefront shaping with an acousto-optic lens for laser scanning microscopy,” Opt. Express 24, 6283 (2016).

22. W. Akemann, S. Wolf, V. Villette, et al., “Fast optical recording of neuronal activity by three-dimensional custom-access serial holography,” Nat. Methods 19, 100–110 (2022).

23. S. Hamann, A. Ceballos, J. Landry, and O. Solgaard, “High-speed random access optical scanning using a linear MEMS phased array,” Opt. Lett. 43, 5455 (2018).

24. O. Tzang, E. Niv, S. Singh, et al., “Wavefront shaping in complex media with a 350 kHz modulator via a 1D-to-2D transform,” Nat. Photonics 13, 788–793 (2019).

25. J. R. Landry, S. S. Hamann, and O. Solgaard, “Random Access Cylindrical Lensing and Beam Steering Using a High-Speed Linear Phased Array,” IEEE Photonics Technol. Lett. 32, 859–862 (2020).

26. P. T. Galwaduge, S. H. Kim, L. E. Grosberg, and E. M. C. Hillman, “Simple wavefront correction framework for two-photon microscopy of in-vivo brain,” Biomed. Opt. Express 6, 2997 (2015).

27. T. Fernández-Alfonso, K. N. S. Nadella, M. F. Iacaruso, et al., “Monitoring synaptic and neuronal activity in 3D with synthetic and genetic indicators using a compact acousto-optic lens two-photon microscope,” J. Neurosci. Methods 222, 69–81 (2014).

28. A. Aggarwal, R. Liu, Y. Chen, et al., “Glutamate indicators with improved activation kinetics and localization for imaging synaptic transmission,” Nat. Methods 20, 925–934 (2023).

29. J.-L. Wu, Y.-Q. Xu, J.-J. Xu, et al., “Ultrafast laser-scanning time-stretch imaging at visible wavelengths,” Light. Sci. & Appl. 6, e16196–e16196 (2016).

30. J. Wu, Y. Liang, S. Chen, et al., “Kilohertz two-photon fluorescence microscopy imaging of neural activity in vivo,” Nat. Methods 17, 287–290 (2020).

31. T. Zhang, O. Hernandez, R. Chrapkiewicz, et al., “Kilohertz two-photon brain imaging in awake mice,” Nat. Methods 16, 1119–1122 (2019).

32. S. Weisenburger, F. Tejera, J. Demas, et al., “Volumetric Ca2+ Imaging in the Mouse Brain Using Hybrid Multiplexed Sculpted Light Microscopy,” Cell 177, 1050–1066.e14 (2019).

33. S. Xiao, I. Davison, and J. Mertz, “Scan multiplier unit for ultrafast laser scanning beyond the inertia limit,” Optica 8, 1403 (2021).

34. A. Kazemipour, O. Novak, D. Flickinger, et al., “Kilohertz frame-rate two-photon tomography,” Nat. Methods 16, 778–786 (2019).

35. A. Shibukawa, R. Higuchi, G. Song, et al., “Large-volume focus control at 10 MHz refresh rate via fast line-scanning amplitude-encoded scattering-assisted holography,” Nat. Commun. 15, 2926 (2024).

36. I. M. Vellekoop, “Controlling the propagation of light in disordered scattering media,” Ph.D. thesis (2008).

37. M. Duocastella, S. Surdo, A. Zunino, et al., “Acousto-optic systems for advanced microscopy,” J. Physics: Photonics 3, 012004 (2021).

38. V. A. Griffiths, A. M. Valera, J. Y. Lau, et al., “Real-time 3D movement correction for two-photon imaging in behaving animals,” Nat. Methods 17, 741–748 (2020).

