## Supplement for "High-speed three-dimensional random access scanning with SPARCLS"

### 1. MICROSCOPE

The microscope setup consists of a 80 MHz femtosecond pulsed laser with included dispersion compensation (MaiTai DeepSee, Spectra-Physics), the scanning module as described in the main text with a f = 32 mm acylindrical lens (AYL4532-B, Thorlabs) and two elliptical mirrors (PFE05-P01, Thorlabs) mounted in a hollow roof prism arrangement, the objective, a detection path and a widefield path for convenient FOV selection.

The laser beam is expanded to 10 mm diameter using a 5x beam expander (GBE05-B, Thorlabs), which enters the scanning module. The SLM plane of the scanning module is then imaged onto the back focal plane of the objective with unit magnification using two achromatic lenses as scan and tube lens (AC508-200-B-ML, Thorlabs). A dichroic mirror (DMLP650L, Thorlabs) separates excitation and emission light. After a bandpass filter (BrightLine 525/50, Semrock), light is detected by a PMT (H10770PA-40 MOD, Hamamatsu), amplified (DHPCA-100, Femto) and detected using a data acquisition card (NI USB-6363, National Instruments). Scanning and data acquisition are synchronized at up to 340 kHz refresh rate and 2 MHz analog input rate using a digital output trigger. The microscope is controlled with software written in Python. The scanning module was designed for a wavelength of 940 nm, optimized for the GFP-based glutamate indicator iGluSnFR3, but can be used at wavelengths in the range of 700-1000 nm with minor changes of alignment.

### 2. SLM CALIBRATION

The SLM pixels consist of ribbons that can be electrostatically actuated by an electrode [1], resulting in a phase modulation of the reflected light equivalent to twice the ribbon deflection, i.e. a $2\pi$ phase delay for a deflection of half the wavelength. Similar to most phase modulators, the use of phase wrapping necessitates the use of (approximately) monochromatic light.

The electrode voltage is proportional to a 10-bit pixel value. As the deflection does not scale linearly with the applied voltage, this needs to be calibrated. Binary gratings with all possible pixel values are applied by varying the pixel value of every other pixel. The deflection d as a function of voltage V can be approximated by $d(V) = AV^4$, where A is a scalar constant [2], which we experimentally determined to be 690.

Calibration shows a maximum stroke of $0.77\pi$ at 940 nm, which reduces the diffraction efficiency only by around 50% (Supplementary Figure S2). Phase patterns (from 0 to $2\pi$) are set to the closest accessible value, i.e. symmetrically clipped at the low and high phase values: As absolute phase does not matter, the range $(1 \pm 0.77/2)\pi$, i.e. ($0.62\pi$ to $1.39\pi$) is displayed in the range from 0 to $0.77\pi$. All values below are set to 0, while values above are set to the maximum, $0.77\pi$.

### 3. AO IMPLEMENTATION

To determine the system correction pattern, we use a modal adaptive optics approach with approximated Zernike polynomials. We iterate through all Zernike polynomials which can be approximated by linearly separable patterns in X and Y with Noll index up to 14. For each Zernike polynomial, the normalized pattern is applied on the phase modulator, multiplied by factors in the range from -5 to 5 in increments of 0.2. Fluorescence intensity of 1 µm beads or bead clusters is determined at 1 kHz SLM refresh rate and the pattern at maximum intensity is determined. This pattern is kept constant while the next mode is added to it. After all modes have been optimized once, we perform a second iteration: the correction pattern determined in the first iteration is applied and only a single mode removed, which is then re-optimized.

### 4. SAMPLE PREPARATION

#### A. Organotypic slice culture preparation

350 mm-thick organotypic hippocampal slice cultures were prepared from P6 to P9 WT C57 mice of either sex. Slices were prepared on filter paper according to the interface method [3, 4] and cultured in a MEM-based mouse slice culture medium, with the addition of 15% Horse Serum; 1x B27; 25 mM HEPES; 3mM L-Glutamine; 2.8 mM CaCl2; 1.8 mM MgSO4; 0.25 mM Ascorbic Acid; 6.5 g/L D-Glucose, adjusted at pH 7.3. Three days after plating, the medium was replaced and then exchanged every 4 days. Cultures were maintained in an incubator with 5% CO2 at 34°C. Plasmids pAAV.hSyn-iGluSnFR3.v857.GPI (Addgene #178331) and pAAV.hSyn.iGluSnFR3.v857.SGZ (Addgene #178330) were gifts from Kaspar Podgorski. The Charité Viral Core Facility manufactured adeno-associated viruses (AAVs). Organotypic hippocampal slice cultures were infected with AAVs at 7-10 days slice culture. Each construct was mixed with mouse slice culture medium to reach 20 μl final volume and pipetted directly on top of the slices. Two photon imaging was performed at 16-23 days slice culture in artificial cerebrospinal fluid (aCSF: 145mM NaCl, 2.5 mM KCl, 10 mM HEPES, 1 mM MgCl2, 2 mM CaCl2, 10 mM glucose; pH 7.3), containing 10 mM KCl and 0.2 mM 4-Aminopyridine (4-AP).

#### B. Zebrafish

iGluSnFR3.GPI was transiently expressed by injecting approximately 20 pg HuC:iGluSnFR3.GPI plasmid DNA into one cell stage zebrafish casper embryos (roy-/-; nacre-/-). Larvae were screened for iGluSnFR expression at 3 days post fertilization (dpf), paralyzed by immersion in 1 mg/mL $\alpha$-bungarotoxin for 2-4 minutes and embedded in 1.5% low melting point agarose. To increase overall activity, pentylenetetrazol (PTZ) was added at a final concentration of 20 mM at least 10 minutes prior to starting an imaging session.

All animal experiments conformed to Berlin state, German federal and European Union animal welfare regulations.

### 5. TRACE ANALYSIS

The baseline fluorescence F0 was assumed to be constant over the measurement time and approximated as the median over the entire trace. dF/F was then computed as dF/F = (F(t)-F0)/F0.

### REFERENCES


1. S. Hamann, A. Ceballos, J. Landry, and O. Solgaard, "High-speed random access optical scanning using a linear MEMS phased array," Opt. Lett. **43**, 5455 (2018).
2. "Silicon Light Machines » IV Response," https://www.siliconlight.com/en/technology/iv-response.html.
3. L. Stoppini, P. A. Buchs, and D. Muller, "A simple method for organotypic cultures of nervous tissue," J. Neurosci. Methods **37**, 173–182 (1991).
4. A. De Simoni and L. MY Yu, "Preparation of organotypic hippocampal slice cultures: Interface method," Nat. Protoc. **1**, 1439–1445 (2006).
5. W. Akemann, J.-F. Léger, C. Ventalon, *et al.*, "Fast spatial beam shaping by acousto-optic diffraction for 3D non-linear microscopy," Opt. Express **23**, 28191 (2015).
6. O. Tzang, E. Niv, S. Singh, *et al.*, "Wavefront shaping in complex media with a 350 kHz modulator via a 1D-to-2D transform," Nat. Photonics **13**, 788–793 (2019).

### 6. SUPPLEMENTARY FIGURES

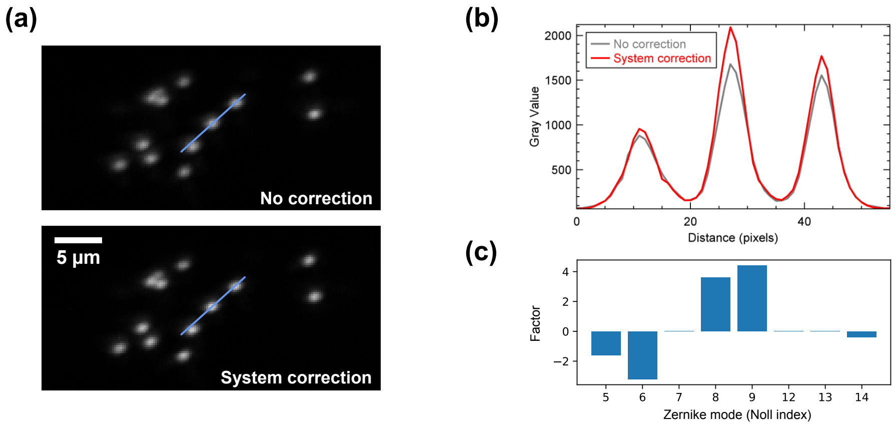


**Fig. S1. Adaptive optics for system correction.** (a) Raster-scanned overview of 1 µm beads (yellow-green FluoSpheres, ThermoFisher) dispersed in agarose. Images are a maximum intensity projection of 9 planes spaced 1 µm apart to minimize the effects of a slight shift in the focal plane. (b) Profiles along the lines shown in (a), showing an increase in signal after system correction (red curve). (c) Contributions of individual Zernike polynomials to the system correction, which can be approximated by a linearly separable pattern. Numbers correspond to Noll indices: 5 - vertical astigmatism, 6 - vertical trefoil, 7 - vertical coma, 8 - horizontal coma, 9 - oblique trefoil, 12 - 1st spherical aberration, 13 - vertical 2nd astigmatism, 14 - vertical quadrafoil. Equations and approximations can be found in Akemann *et al.* [5].

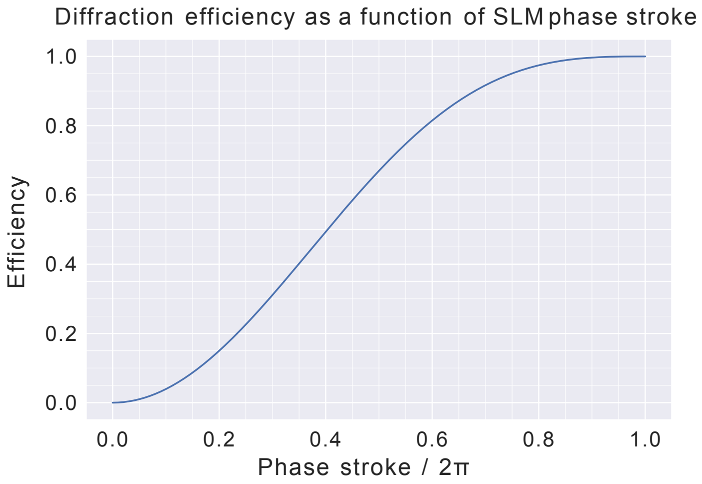


**Fig. S2. Diffraction efficiency as a function of maximum phase stroke.** Numerical simulation assumes a target pattern (focused spot), which is Fourier transformed, the phase range clipped and then transformed back. The efficiency is the energy in the new target pattern normalized to the old. See also Tzang *et al.* [6].

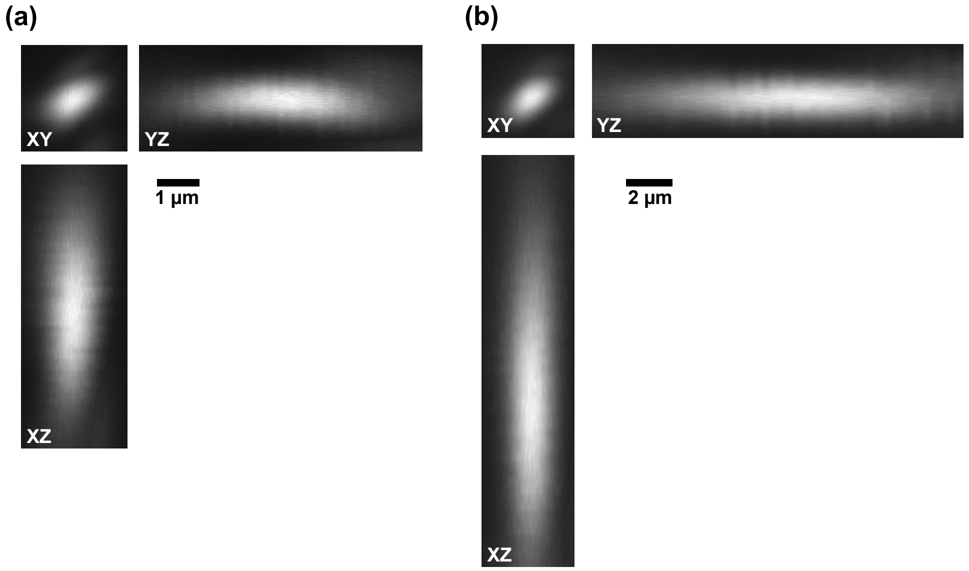


**Fig. S3. Point spread functions.** Maximum intensity projections in the XY, XZ and YZ planes using (a) a 40x/0.8NA objective (FWHM x: 1.0 μm, y: 0.8 μm, z: 3.5 μm) and (b) a 25x objective with effective illumination NA of 0.63 due to underfilling (FWHM x: 1.6 μm, y: 1.3 μm, z: 9.3 μm). The lateral PSF size represents an upper bound, as we used 1 μm diameter beads. The PSF is not diffraction-limited. This is due to aberrations which are not linearly separable, and thus cannot be corrected by the scanning module, as well as chromatic aberration due to the acylindrical lens. In the future, this could be improved by a custom achromatic cylindrical instead of an off-the shelf acylindrical lens.

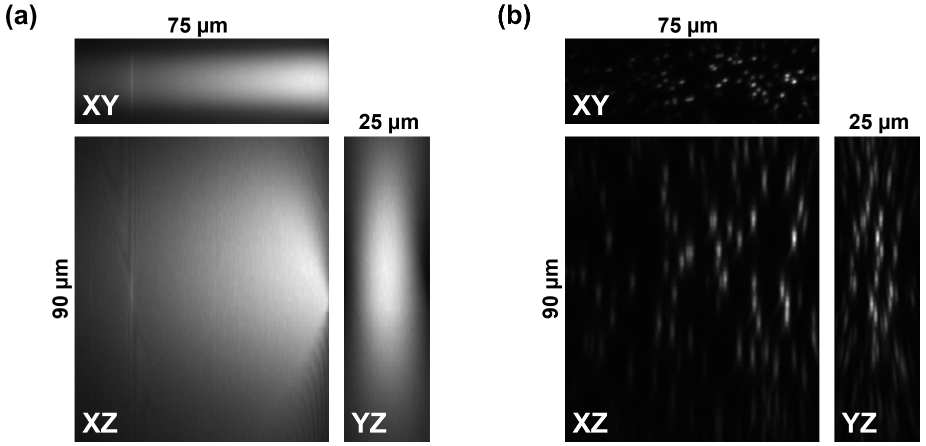


**Fig. S4. Field of view using the 40x/0.8NA objective.** (a) Maximum intensity projections along all three axes visualized using a fluorescein solution. (b) FOV visualization using 1 μm diameter beads (yellow-green FluoSpheres, ThermoFisher).

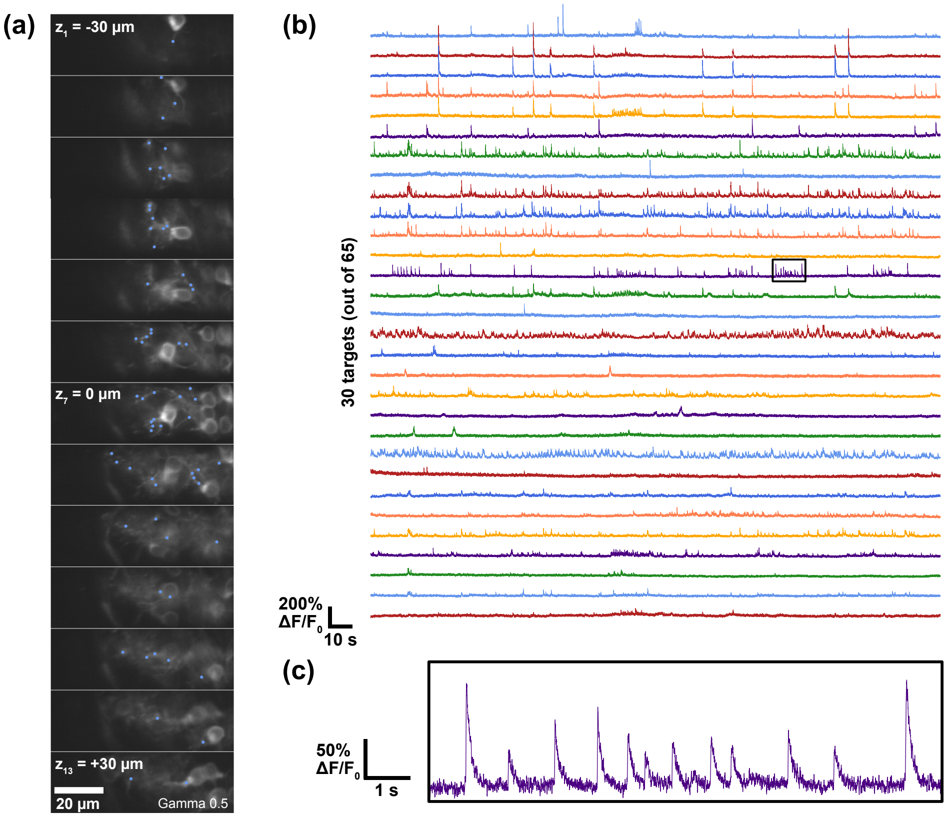


**Fig. S5. 3D random access microscopy using iGluSnFR3 in larval zebrafish olfactory bulb at 100 kHz point rate.** (a) Raster scanned overview stack of part of the olfactory bulb of a 4 dpf zebrafish larva. A Gamma correction was applied to emphasize dimmer structures. Manually selected targets for random-access scanning are marked as blue dots. (b) Change in fluorescence dF/F0 acquired at 100 kHz. A 20 point moving average is applied. (c) Zoom in of the region in (b) marked with a black rectangle.
